# Dichotomy of neutralizing antibody, B cell and T cell responses to SARS-CoV-2 vaccination and protection in healthy adults

**DOI:** 10.1101/2023.05.24.541920

**Authors:** Edward J Carr, Hermaleigh Townsley, Mary Y Wu, Katalin A Wilkinson, Philip S Hobson, Dina Levi, Sina Namjou, Harriet V Mears, Agnieszka Hobbs, Martina Ragno, Lou S Herman, Ruth Harvey, Chris Bailey, Ashley S Fowler, Emine Hatipoglu, Yenting Ngai, Bobbi Clayton, Murad Miah, Philip Bawumia, Mauro Miranda, Callie Smith, Chelsea Sawyer, Gavin Kelly, Viyaasan Mahalingasivam, Bang Zheng, Stephen JW Evans, Vincenzo Libri, Andrew Riddell, Jerome Nicod, Nicola O’Reilly, Michael Howell, Bryan Williams, Robert J Wilkinson, George Kassiotis, Charles Swanton, Sonia Gandhi, Rupert CL Beale, David LV Bauer, Emma C Wall

**Affiliations:** The Francis Crick Institute, 1 Midland Road, London, NW1 1AT, UK; National Institute for Health Research (NIHR) University College London Hospitals (UCLH) Biomedical Research Centre and NIHR UCLH Clinical Research Facility, London, UK; Covid Surveillance Unit, The Francis Crick Institute, 1 Midland Road, London, NW1 1AT, UK; Wellcome Centre for infectious Diseases Research in Africa, University of Cape Town, Observatory, South Africa; Worldwide Influenza Centre, The Francis Crick Institute, 1 Midland Road, London, NW1 1AT, UK; London School of Hygiene & Tropical Medicine, London, WC1E 7HT, UK; University College London, Gower Street, London, UK; Imperial College London, SW7 2AZ, UK; Department of Infectious Disease, St Mary’s Hospital, Imperial College London, London, W12 0NN, UK; Genotype-to-Phenotype UK National Virology Consortium (G2P-UK)

## Abstract

Heterogeneity in SARS-CoV-2 vaccine responses is not understood. Here, we identify four patterns of live-virus neutralizing antibody responses: individuals with hybrid immunity (with confirmed prior infection); rare individuals with low responses (paucity of S1-binding antibodies); and surprisingly, two further groups with distinct serological repertoires. One group – broad responders – neutralize a range of SARS-CoV-2 variants, whereas the other – narrow responders – neutralize fewer, less divergent variants. This heterogeneity does not correlate with Ancestral S1-binding antibody, rather the quality of the serological response. Furthermore, IgD^low^CD27^-^CD137^+^ B cells and CCR6^+^ CD4^+^ T cells are enriched in broad responders before dose 3. Notably, broad responders have significantly longer infection-free time after their third dose. Understanding the control and persistence of these serological profiles could allow personalized approaches to enhance serological breadth after vaccination.

## Introduction

Inter-individual heterogeneity in human immune responses is well-described ^1–4^. While immunological heterogeneity was previously seen as a “nuisance variable”, over the last 10-20 years, this view has shifted due to new strategies, tools and hypotheses. Today, the study of the variation between individuals provides novel insights in human immunology ^5^.

Alongside the deployment of COVID-19 vaccines as the primary control strategy of the SARS-CoV-2 pandemic, observational studies were established to generate data to inform timing of future doses, and to examine vaccine immunogenicity in vulnerable populations omitted from phase 3 trials. Secondary aims included studying the mechanisms of vaccine responses for mRNA and adenoviral vectored vaccines, vaccine platforms not previously used outside of early phase trials, with profound primary immunological and epidemiological responses ^6–8^. We, with colleagues, established three sentinel UK studies: CAPTURE, studying responses in solid-organ and hematological cancer patients ^9–11^, NAOMI exploring responses in hemodialysis patients ^12, 13^, and the Legacy study, an observational cohort study of healthy adults undergoing occupational health screening and vaccination for SARS-CoV-2 ^14–16^. Legacy is a collaboration between the Francis Crick Institute and University College London Hospitals (clinical trials registration NCT04750356). We have previously reported on neutralizing ability of sera after two ^14, 15^, and three doses of vaccine ^16^. Here, we explore inter-individual differences in serological and cellular responses before and after a third vaccine dose in 283 Legacy participants (**Fig. 1A**) and show that stratification of individuals based on live-virus neutralization patterns uncovers previously unrecognized immune differences in otherwise healthy individuals.

**Figure 1.**
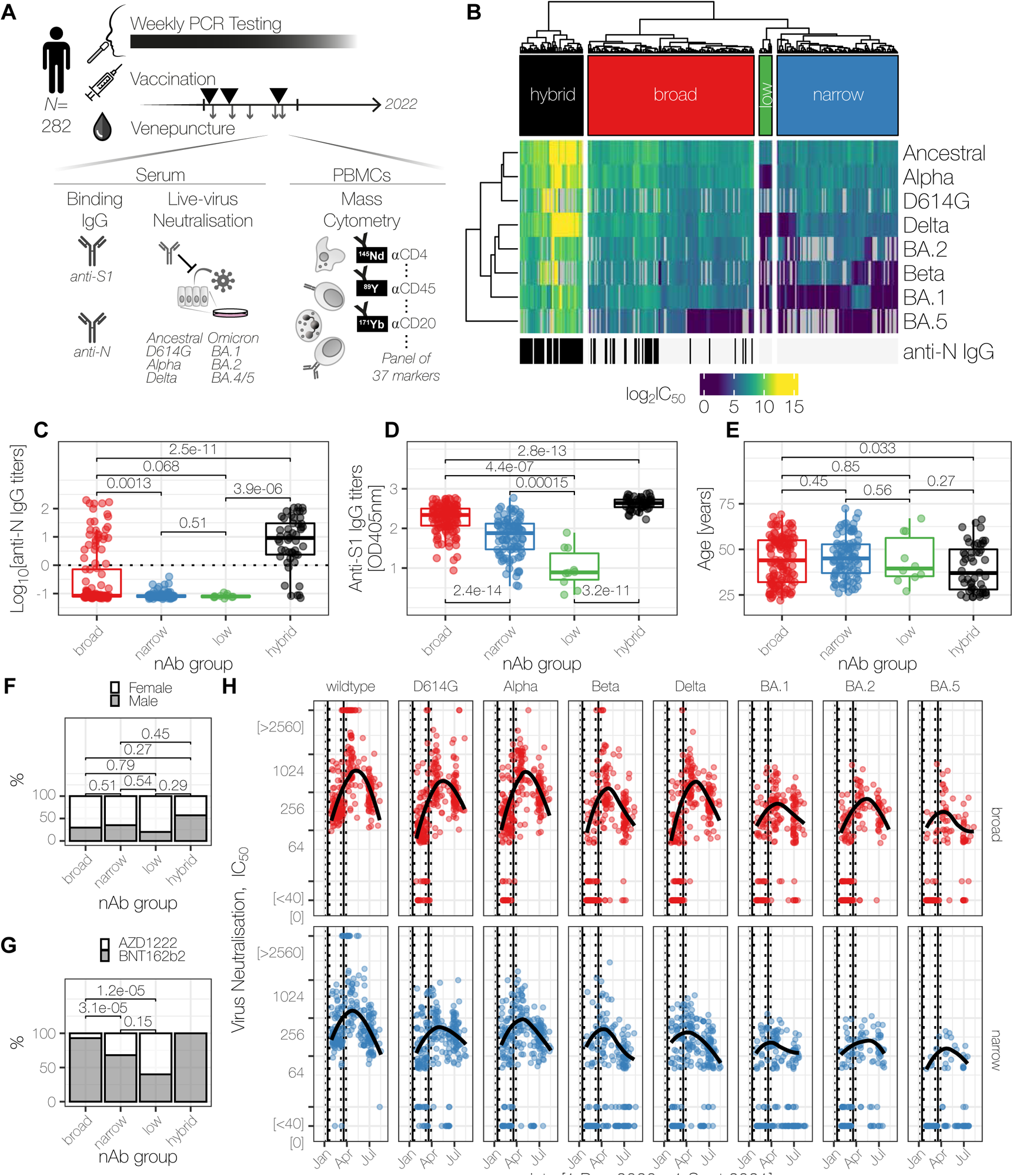
Serological profiling of Legacy participants identifies heterogeneity in patterns of SARS-CoV-2 neutralization. **(A)** Study design, longitudinal sampling and assays performed for 282 healthcare or laboratory workers. **(B)** Hierarchical clustering of live-virus neutralizing antibody titers before dose 3. Each individual is represented by a column and each SARS-CoV-2 variant by a row. The log_2_IC_50_ is shown by the color bar, and missing data are in grey. Both rows and columns are clustered using Euclidean distances and anti-N IgG status is indicated. In the color bar above the heatmap, the label of each groups is shown: hybrid, broad, low, and narrow responders in black, red, green and blue respectively. **(C)** anti-N IgG titers. **(D)** anti-S1 IgG titers. **(E)** participant age at enrollment. **(F)** Participant sex, and **(G)** vaccines used for doses 1 and 2, for anti-N negative individuals. **(H)** Trajectory neutralizing antibody titers between doses 1 and 3 of anti-N negative individuals from broad and narrow clusters. In (C)-(E) and (F)-(G) *P* values are from two-tailed unpaired Mann-Whitney tests or χ^2^ tests respectively. In (H), smoothed splines were restricted to data within the quantitative range of the assay, and vertical solid and dashed lines represent the median and inter-quartile ranges of the dates of doses 1 and 2.

## Results

We hypothesized that inter-individual heterogeneity would exist in the neutralizing antibody responses after SARS-CoV-2 vaccination. We anticipated identifying three groups: firstly, those with strong neutralization capacity due to encounters with Spike during infection episodes, in addition to their vaccinations – so called “hybrid” immunity ^17^; secondly, a small number of partial or low responders where their vaccine-induced antibody responses were attenuated; and thirdly, a group of “normal” responders, comprising the bulk of participants. To test this hypothesis, we performed hierarchical clustering of the neutralizing capacity of 282 pre-dose 3 sera against ancestral SARS-CoV-2 and seven variants of concern (VOCs, **Fig. 1B**). The first two doses were either AZD1222 (Oxford/AstraZeneca, n=73) or BNT162b2 (Pfizer-BioNTech, n=209). Surprisingly, unsupervised clustering identified four groups of individuals, which we tentatively assigned: individuals with hybrid responses (n=49, 17%), “low responders” (n=10, 3.5%) and two unexpected further groups, “broad responders” (n=129, 46%) and “narrow responders” (n=94, 33%), mainly separated by their neutralization (or not) of Omicron BA.1 before dose 3. To confirm the biological identities of these clusters, we assessed whether we could identify these four groups using infection history, anti-S1 and anti-N IgG. We proposed that hybrid responses should be readily identifiable with exposure history and widely available binding S antibody and anti-N IgG assays. This strategy confirmed our grouping of individuals with hybrid immunity: 46 individuals (of 49; 94%) had 47 episodes of prior infections confirmed by the presence of symptoms (39 episodes, 87.8%), by a molecular test (31 episodes, 66%), or by the detection of anti-nucleocapsid IgG (44 individuals, 93.6%) (**Fig. 1C**). The ten low responders were separable from the rest of the cohort by low anti-S1 IgG titers (**Fig. 1D**), and neutralizing activity restricted to ancestral SARS-CoV-2 (**Fig. 1B**). Hybrid and low responders could be identified by binding anti-S1 and anti-N IgG individually or jointly, however these tests distinguished poorly between broad and narrow responders (**Fig. S1**).

Having confirmed two biologically relevant groups – low and hybrid responders – through exposure history and anti-S1/anti-N titers, we next focused on the broad and narrow responder groups which were not clearly defined by these parameters. We reasoned that these two groups were likely to reflect similarly important, but hitherto unrecognized, biological distinctions and therefore sought to further characterize these groups. From hierarchical clustering with serum drawn just before dose 3, we observed that 119 of 129 [92.3%] broad responders had serum IC_50_>40 for Omicron BA.1, indicating neutralizing activity against the Omicron BA.1 lineage, whereas only 19 of 94 [20.3%] narrow responders had serum IC_50_>40 for Omicron BA.1 (χ^2^ test *P*<2.2×10^-16^; **Fig. 1B**). Neutralization titers against Omicron BA.1 before dose 3 therefore offer a population-level surrogate. We found a small fraction of the broad group was also anti-N positive (31 of 130, 23.8%; **Fig. 1C**), from prior infection. The vast majority of broad responders was not previously infected by SARS-CoV-2, based on weekly occupational health screening by RT-qPCR for SARS-CoV-2 infection and absence of anti-N IgG (**Fig. S2**). Anti-N IgG positive broad responders were positive from their first serum sample, indicating infection in 2020 **(Fig. S2C)**. Focusing on anti-N IgG negative individuals, there were no differences in age and sex between broad and narrow responders (**Fig. 1E-F**). We found that BNT162b2 was more commonly used for doses 1 and 2 than AZD1222 in broad responders, compared to either narrow or low responders (**Fig. 1G**, χ^2^ test *P*= 0.006 or 0.002 respectively). We excluded benign explanations for the difference between these two groups: there were no differences in age or gender between broad N seronegative (N-) or seropositive individuals (**Fig. S3A-B**); broad N-individuals were more likely to have been administered BNT162b2 for doses 1 and 2 (**Fig. S3C**); additional spike exposure through infection provided boosting to anti-S1 titers in broad N+ individuals compared to N-individuals (**Fig. S3D**); and there were no differences between any of the four groups in intervals between doses 1 and 2 or between dose 2 and their serum sample (**Fig. S3E-F**).

Interestingly, we plotted the trajectories of neutralizing titers against 8 different variants between doses 2 and 3, and found, that broad and narrow responder groups followed offset trajectories throughout this period, suggesting that an individual’s response is consistently either broad or narrow, across antigen encounters (**Fig. 1H**). Next, we considered whether serological breadth initiated by SARS-CoV-2 would include other coronaviruses. To test this possibility, we performed live-virus microneutralization assays using HCoV-OC43, a seasonal human coronavirus (**Fig. S4**). We found no differences in starting titers between broad or narrow responders, and no boosting effect from SARS-CoV-2 vaccination in either group.

Given that neutralizing antibody production is a function of the orchestrated response of B and CD4^+^ T cells after vaccination, we anticipated that underlying lymphocyte differentiation might give rise to our observed distinct serological profiles. To determine whether cellular differences contribute to neutralization breadth we performed mass cytometry in individuals with broad or narrow serological profiles (n=11 and 6 respectively), before (median 1d [range 10-0]) and after (median 19d [range 14-21]) their third BNT162b2 dose (**Table S1**). All individuals were anti-N IgG negative at both timepoints, and all preceding samples. Gating, quality control and clustering are described in the Methods (**Fig. S5-7**). First, we assessed whether changes in the B cell compartment were present between individuals with broad and narrow serological profiles (**Fig. 2A and B**). We expected altered utilization of different memory B cell compartments between broad and narrow responders. We therefore compared the pre-dose 3 samples (*i.e.* the long-term memory footprint from dose 2) between broad and narrow responders, and found that IgD^low^CD27^-^CD137^+^ B cells were more abundant in the broad responders (cluster B3: log_2_ fold change 4.3, *P*_adj_ 0.045 and cluster B7: log_2_ fold change 4.1, *P*_adj_ 0.013, **Fig. 2C**). IgD^low^CD27^-^ B cells are traditionally termed double negative (DN) memory cells, originally described in ageing and chronic infections and now with newer evidence from many groups showing roles in healthy serological responses (reviewed in ^18^). Following dose 3, there were no differences in the B cell compartment between broad and narrow responders that reached statistical significance comparing before and after third-dose vaccination (**Fig. 2D**).

**Figure 2.**
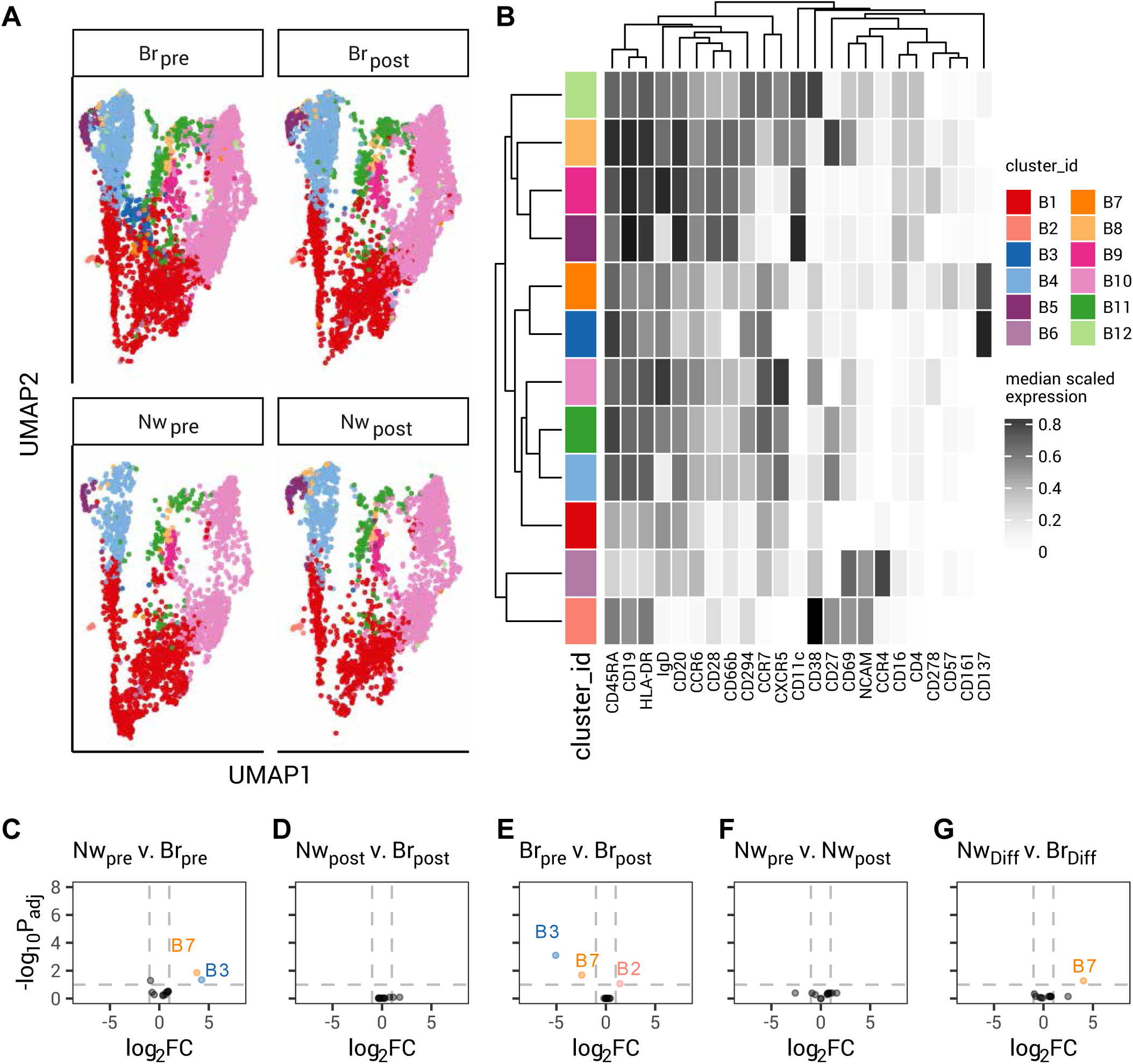
Mass cytometry demonstrates altered B cell sub-populations between broad and narrow responders before and after third doses. **(A)** UMAP embedding of B cells separated by breadth (Nw narrow, n=6; Br broad, n=11) and before (pre) and after (post) vaccination. 12 clusters identified by FlowSOM and ConsensusClusteringPlus are shaded. (**B)** Heatmap of surface expression of selected markers for the clusters shown in (A). Rows represent the clusters shown in (A), and their color key is shared. Columns reflect the labelled cell surface marker. Scaled expression is shown from white (low/no expression) to black (high expression). **(C)**-**(G)** Differential abundance analysis for the 12 B cell clusters shown in (A) and (B), for the comparisons indicated: narrow pre vs broad pre in (C); narrow post vs broad post in (D); broad pre vs.broad post in (E); narrow pre vs.narrow post in (F); and the difference between (narrow pre vs. narrow post) and (broad pre vs. broad post) in (G). For (C-G), log_2_ fold change ±1 and *P*_adj_=0.01 are shown by dashed lines. Color keys are shared (A-G).

The next cellular comparison was the response to vaccination within each serological profile. Comparing broad responders before and after vaccination, we found a decrease in the abundance of atypical, double-negative memory B cells that express CD137 (IgD^low^CD27^-^CD137^+^; cluster B3: log_2_ fold change -5.1, *P*_adj_ 0.0008 and cluster B7: log_2_ fold change -2.4, *P*_adj_ 0.02, **Fig. 2E**). In narrow responders, we found no changes in the B cell compartment after vaccination (**Fig. 2F**). For plasmablasts (cluster B2: CD20^-^CD27^+^CD38^+++^), we observed an expansion in broad responders, which did not reach our significance threshold (log_2_ fold change 1.4, *P*_adj_ 0.08) and a smaller, non-significant fold-change in narrow responders (log_2_ fold change 0.6, *P*_uncorrected_ 0.42, *P*_adj_ 0.52). Finally, we tested for differential vaccine responses between each serological profile, and found no significant differences (**Fig. 2G,** cluster B7 *P*_adj_=0.053).

Together, these results suggest that broad responders favor a relatively higher proportion of DN-CD137^+^ memory B cells after two doses, which is perturbed by further vaccination. Broad responders also had a tendency towards larger plasmablast responses after dose 3. CD137 expression on human B cells has been shown in several contexts, including CD11c^+^ B cells (a further subgroup of DN memory cells) in healthy donors, lupus, and systemic sclerosis ^19^; healthy B cells stimulated *in vitro* ^20^; and lymphoma, including on Hodgkin Reed-Sternberg cells ^20, 21^. There are reports of rare individuals with de-functioning mutations in *TNFRSF9*, the gene encoding CD137 (also called 4-1BB), who display perturbations in B cell biology including a propensity to autoinflammation and lymphomagenesis, vulnerability to respiratory infections, and attenuated responses to vaccination ^22, 23^.

Since B cell memory development is cued in part by CD4^+^ T_H_ cells, we next assessed the CD4^+^ T_H_ cell compartment (**Fig. 3A-B**). Before third doses, we found two clusters were over-represented in narrow responders (cluster H8 CXCR3^+^TCRψο^+^: log_2_ fold change -5.6, P_adj_ 3.1 x 10^-6^; cluster H11 NCAM^+^CXCR3^+^TCRψο^+^: log_2_ fold change - 4.3, P_adj_ 5.9×10^-4^, **Fig. 3C**), and found a CCR7^-^CD27^-^CD28^-^CD45RA^+^CCR6^+^CD57^+^ population (cluster H12) that was “pre-expanded” in broad responders (log_2_ fold change 4.5 and P_adj_ 0.01, **Fig. 3C**). For brevity, we refer to these CD27^-^CD28^-^ CD45RA^+^CCR6^+^CD57^+^ CD4^+^ T cells as breadth-related T_H_ cells (brT_H_). We found no significant differences comparing CD4^+^ T cells between the two serological profiles after vaccination (**Fig. 3D**). Comparing before and after third doses, we found no significant differences in the abundance of CD4^+^ T_H_ cell clusters in broad responders (**Fig. 3E**). However, narrow responders showed a significant decrease in two clusters in response to vaccination (cluster H8: log_2_ fold change -6.3, P_adj_ 0.0002; cluster H11: log_2_ fold change -6.4, P_adj_ 0.0002, **Fig. 3F**). CD4^+^T cells co-expressing TCRψ8^+^ and TCRαβ^+^ have been recently described ^24^. Clusters H8 and H11 were distinct from *bona fide* TCRψ8^+^ T cells, which we observed as a distinct population adjacent to CD8 T cells in our PBMC analysis (**Fig. S6**). The only differentially responsive clusters between broad and narrow individuals were clusters 8 and 11 (**Fig. 3G**). brT_H_ are CCR7^-^CD45RA^+^ suggesting a terminally differentiated effector memory phenotype (T_emra_) ^25^, and further classifiable within a CD27-CD28-T_emra_ sub-compartment ^26^. A study examining CD4 responses to Dengue virus, reported two subgroups of Dengue-specific CD4^+^ T_emra_, one of which was CCR6^+^ and lacked the expression of cytotoxic and terminal differentiation markers (perforin and KLRG1) found on CCR6^-^ T_emra_ ^27^. Taken together, these observations suggest that broad responders harbor T_emra_-like memory populations, which are not terminally differentiated.

**Figure 3.**
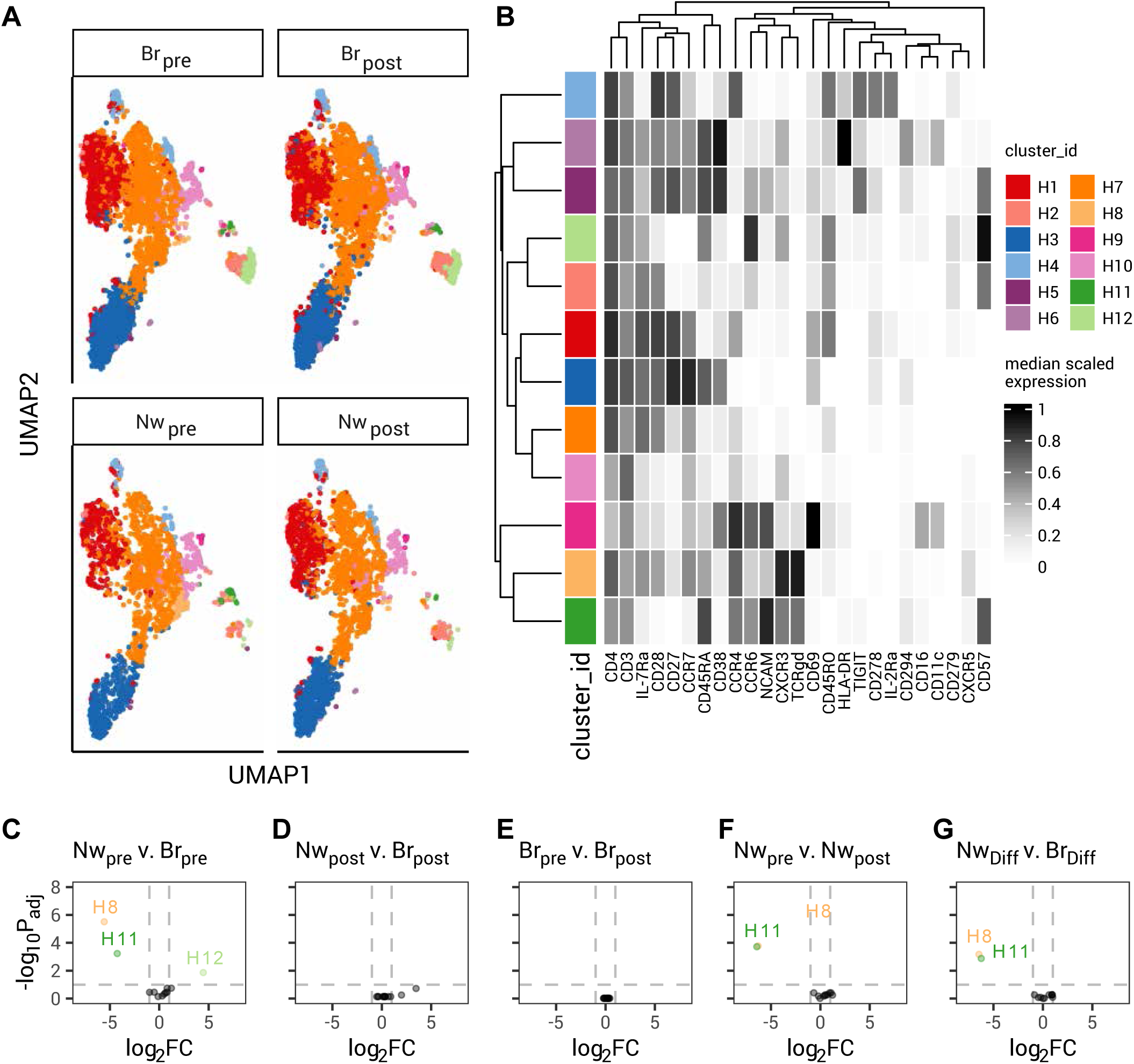
Perturbations in the CD4^+^ T cell compartment between broad and narrow responders before and after third doses. **(A)** UMAP embedding of CD4^+^ T cells separated by breadth (Nw narrow, n=6; Br broad, n=11) and before (pre) and after (post) vaccination. 12 clusters identified by FlowSOM and ConsensusClusteringPlus are shaded. (**B)** Heatmap of surface expression of selected markers for the clusters shown in (A). Rows represent the clusters shown in (A), and their color key is shared. Columns reflect the labelled cell surface marker. Scaled expression is shown from white (low/no expression) to black (high expression). **(C)**-**(G)** Differential abundance analysis for the 12 CD4^+^ T cell clusters shown in (A) and (B), for the comparisons indicated: narrow pre vs broad pre in (C); narrow post vs broad post in (D); broad pre vs.broad post in (E); narrow pre vs.narrow post in (E); and the difference between (narrow pre vs. narrow post) and (broad pre vs. broad post) in (G). For (C-G), log_2_ fold change ±1 and *P*_adj_=0.01 are shown by dashed lines. Color keys are shared (A-G).

To summarize our multi-dimensional cytometry: examining differences between individuals with broad and narrow serological profiles showed that broad responders are marked by the presence of brT_H_ cells and DN-CD137^+^ B cells before their third doses. We found a propensity for broad responders to expand their plasmablast population compared to narrow responders. After the immune stimulus of a third mRNA vaccination, there were no cellular differences in either B or CD4^+^T_H_ cells populations between broad and narrow responders (**Fig. 2D and 3D**). There are several possible explanations for this observation. Firstly, vaccination has been shown to transiently perturb the immune landscape ^1, 2^, thus cytometry performed during a time of immune activation may be obfuscated, and overlook intrinsic underlying inter-individual differences. In this situation, a later timepoint, memory analysis of post-vaccination may be more informative, as it lacks the overlaid perturbation from a recent vaccination. Our pre-dose 3 samples provide that retrospective memory assessment of dose 2, once the early cellular changes have resolved, and is it at that timepoint that we observed the most striking cellular differences between broad and narrow responders. Secondly, it is possible that narrow responders represent an accelerated resolution of the cellular changes after doses 1 and 2, with the loss of certain memory lymphocyte sub-populations rapidly (brT_H_ cells and DN-CD137^+^ B cells), whereas broad responders retain a diversity in their cellular responses until after dose 3. If a third dose were to “reset” the serological profile such that there are no cellular or serological differences between broad or narrow responders (defined before dose 3), then we would anticipate that all individuals would have equal susceptibility to infection after dose 3, as neutralizing antibody is the single best predictor of infection ^28, 29^.

Thus, we next assessed whether membership of these two serologically-defined groups of broad and narrow responders influenced susceptibility to SARS-CoV-2 infection, after their third doses with Omicron BA.2. Because we expected additional antigenic exposures to influence breadth (**Fig. 1B**), we censored individuals with identified BA.1 infection, or who seroconverted to nucleocapsid (at the date of their first positive anti-N IgG result). Individuals from the lowest age quartile (22-33yo) had a tendency towards an increased likelihood of experiencing an infection compared to those from the highest age quartile (53-72yo; **Fig. 4A**). There were no sex-related differences (**Fig. 4B**). We found that our two serological profiles of interest, broad and narrow, were significant predictors of time-to-infection (**Fig. 4C**), with participants older than the median age (>44yo) from the broad group protected from infection relative to their counterparts in the narrow group. The serological effect was attenuated in 22-44yo (**Fig. 4C**). To quantify the effect of breadth across the entire age range, we fitted a Cox proportional hazard model, allowing interactions between breadth and age, and dividing age into two groups: those older (>44yo) or younger (22-44yo) than the median (**Fig. 4D**). This model gave a hazard ratio for broad responders of 0.45 (HR, 95% CI 0.22-0.94) in the >44yo age group (the reference age group), implying a ∼60% reduction in infection risk during the Omicron BA.2 wave for broad responders compared to narrow responders. The interaction term between age and breadth suggest that the serological effect was attenuated in younger participants approximately two-fold (**Fig. 4D**). A potential limitation of this analysis is that the timing of the BA.2 in the UK was at a time when asymptomatic community and occupational testing was being withdrawn, and when national requirements to isolate after a positive test ceased, so it is possible the exposure risk varied. In summary, we found that serologically defined groups of individuals with altered B and T cell compartments were differentially protected from infection after vaccination, especially among older adults in our cohort.

**Figure 4.**
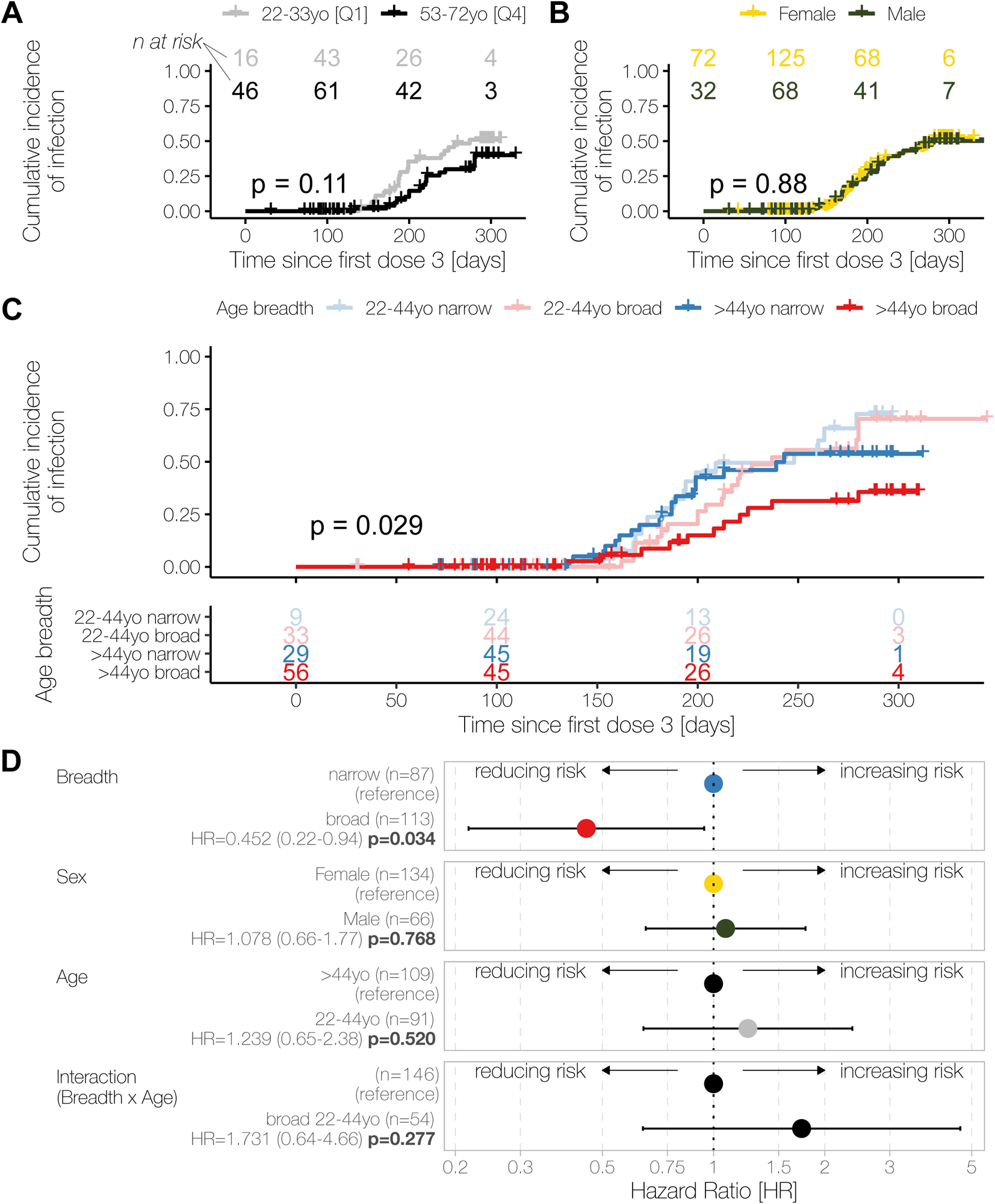
Individuals with broad neutralizing responses are relatively protected from Omicron BA.2 infection. **(A)** and **(B)** Time-to-event analysis for the acquisition of an Omicron BA.2 infection >14d after dose 3 in individuals in the first and last age quartiles: 22-33 years old (yo; Q1, grey) or 53-72 yo (Q4, black) (A); or in female (yellow) or male (green) participants (B). **(C)** Time-to-event analysis for the acquisition of an Omicron BA.2 infection >14d after dose 3 with two age groups (22-44yo, Q1-Q2; 44-72yo,>44yo, Q3-Q4), and by breadth of neutralization responses before dose 3. **(D)** Forest plot of proportional hazard ratios from a Cox proportional hazard model, with breadth, sex, age groups and the interaction term between breadth and age as predictors. For age, >44yo is used as the reference group (hazard ratio, HR=1); for breadth and sex, narrow and female are used as the respective reference group (HR=1). In (A)-(D), left-censoring occurs on the day of that individual’s third dose and right censored with a non-BA.2 infection, or their last study visit before their fourth dose. In (A)-(C), the x-axis is the time in days since the earliest third dose, and P values are the log likelihood ratio test from a Cox model. The numbers at risk are shown for each group within the graph (A) and (B), or tabulated below (C).

## Discussion

Here, we have used detailed serological profiling to uncover inter-individual heterogeneity in vaccine responses, with corresponding alterations in T and B cell compartments, and investigated the relationship between these differences in immunity and subsequent risk of SARS-CoV-2 infection. Serological profiling with live-virus microneutralization assays identified 4 groups – responders with hybrid immunity, and those with low, narrow, or broad responses. We have focused on the apparent dichotomy between the cohort of broad responders who have serological capacity to neutralize Omicron lineages before their third doses, and the cohort of narrow responders who do not. We have shown that surrogate classification by binding anti-S titers by ELISA is inadequate to define these classes of breadth; a range of neutralization titers against a panel of viruses is required. We have found that broad responders have specific lymphocyte populations in circulation before dose 3, including DN-CD137^+^ B cells and brT_H_. Our findings are critical in several contexts. Firstly, to offer personalized risk assessments to current VOCs, or forwards prognostication, anti-S is inadequate. Secondly, the inter-individual heterogeneity appears consistent over ∼60-100 days — without additional antigen encounter from infection: evidenced by symptom diaries, PCR screening, anti-N IgG testing (**Fig. S2**). This observation suggests that breadth might be intrinsic to that individual, with implications for other vaccine responses (and design), and perhaps for antibody responses in general including autoimmune contexts. Thirdly, variant-specific booster trials will require careful interpretation: inadvertently unmatched arms between broad and narrow could plausibly reverse or obfuscate a true effect. In conclusion, we show that our serological profiling with high-throughput live-virus microneutralization identifies immunological and epidemiological inter-individual heterogeneity, where breadth of neutralizing response is key to protection. Our data suggest serological breadth of response to vaccination is not a purely stochastic phenomenon in humans, rather it has important underlying cellular correlates with fertile ground for further study to understand both the mechanistic underpinnings and their clinical consequences.

## Materials and Methods

### Ethics approvals and study design

The Legacy study (NCT04750356) was established in January 2021 and enrolled two prospective cohorts. The Legacy study was approved by London Camden and Kings Cross Health Research Authority (HRA) Research and Ethics committee (REC) IRAS number 286469 and sponsored by University College London, The study has been described in our prior interim reports ^14–16^. Participants were included if they were an employee of either UCLH or the Francis Crick Institute and had provided at least one swab for qRT-PCR testing via the Crick PCR pipeline. At the commencement of the Legacy study, the Crick PCR pipeline was performing NHS staff and patient testing to support local NHS Trusts and partners. Participants comprised of patient facing healthcare workers at UCLH and Crick staff. Study visits with venipuncture were offered approximately one month after vaccination, and at approximately 3, 6 and 12 months. Participants who experienced infection after two (or more) doses of vaccine were invited for a study visit approximately 2 weeks after the start of their infection episode.

### SARS-CoV-2 RT-qPCR

RNA was extracted from nasopharyngeal swabs taken at time of occupational health screening, as previously described ^30^. Viral RNA was detected by RT-qPCR (TaqPath COVID-19 CE-IVD Kit, ThermoFisher) to confirm SARS-CoV-2 infection. Individuals reporting symptoms, positive lateral flow tests, or positive external PCR testing were invited to perform a study nasopharyngeal swab.

### Venipuncture and serum processing

Legacy participants were invited for venipuncture before and after (∼10-21d) vaccinations, with additional samples planned at approximately 3, 6 and 12 months. After an infection episode, individuals were invited for additional venipuncture after convalescence (∼10-21d). Venipuncture was performed into K2-EDTA (for PBMC), or SST (serum) vacutainer tubes (BD). Serum was separated within 24 hours.

### PBMC isolation

Whole blood was collected in K2-EDTA tubes and samples were processed within 24 hours. PBMC and plasma were isolated by density-gradient centrifugation for 30 minutes at 1000 x g at RT. Plasma was carefully removed then centrifuged for 10 minutes at 4000 x g to remove debris, aliquoted and stored at -80°C. The cell layer was then collected and washed twice in PBS by centrifugation for 10 minutes at 300 x g at RT. PBMC were resuspended in cell freezing medium (Fisher Scientific) containing 10% DMSO, placed overnight in CoolCell freezing containers (Corning) at -80°C and then stored in liquid nitrogen tanks until batched analysis.

### Virus variants and culture

The Alpha, Delta and Omicron BA.1 isolates used were the same as previously, and our viral culture technique is unchanged ^14–16^. The SARS-CoV-2 B.1.1.7 isolate (“Alpha”) was hCoV-19/England/204690005/2020, which carries the D614G, Δ69-70, Δ144, N501Y, A570D, P681H, T716I, S982A and D1118H mutations in Spike ^31^, and was obtained from Public Health England (PHE), UK, through Prof. Wendy Barclay, Imperial College London, London, UK via the Genotype-to-Phenotype National Virology Consortium (G2P-UK). The B.1.617.2 (“Delta”) isolate was MS066352H (GISAID accession number EPI_ISL_1731019), which carries the T19R, K77R, G142D, Δ156-157/R158G, A222V, L452R, T478K, D614G, P681R, D950N mutations in Spike, and was kindly provided by Prof. Wendy Barclay, Imperial College London, London, UK via the Genotype-to-Phenotype National Virology Consortium (G2P-UK). The BA.1 (“Omicron”) isolate was M21021166, which carries the A67V, Δ69-70, T95I, Δ142-144, Y145D, Δ211, L212I, G339D, S371L, S373P, S375F, K417N, N440K, G446S, S477N, T478K, E484A, Q493R, G496S, Q498R, N501Y, Y505H, T547K, D614G, H655Y, N679K, P681H, A701V, N764K, D796Y, N856K, Q954H, N969K, and L981F mutations in Spike, and was kindly provided by Prof. Gavin Screaton, University of Oxford, Oxford, UK via G2P-UK. The Omicron BA.2 isolate carries the T19I, Δ24-26, A27S, G142D, V213G, G339D, S371F, S373P, S375F, T376A, D405N, R408S, K417N, N440K, S477N, T478K, E484A, Q493R, Q498R, N501Y, Y505H, D614G, H655Y, N679K, P681H, N764K, D796Y, Q954H, and N969K mutations in Spike and was obtained from a Legacy study participant. The Omicron BA.2.12.1 isolate carries the L452Q and S704L mutations in Spike, in addition to the BA.2 mutations listed previously, and was kindly provided by Prof. Gavin Screaton, University of Oxford, Oxford, UK. The Omicron BA.5 isolate carries the T19I, Δ24-26, A27S, Δ69-70, G142D, V213G, G339D, S371F, S373P, S375F, T376A, D405N, R408S, K417N, N440K, L452R, S477N, T478K, E484A, F486V, Q498R, N501Y, Y505H, D614G, H655Y, N679K, P681H, N764K, D796Y, Q954H, and N969K mutations in Spike was obtained from the laboratory of Alex Sigal, Africa Health Research Institute, Durban, South Africa.

All viral isolates were propagated in Vero V1 cells (a gift from Stephen Goodbourn). Briefly, 50% confluent monolayers of Vero V1 cells were infected with the given SARS CoV-2 strains at an MOI of approx. 0.001. Cells were washed once with DMEM (Sigma; D6429), then 5 ml virus inoculum made up in DMEM was added to each T175 flask and incubated at room temperature for 30 minutes. DMEM + 1% FCS (Biosera; FB-1001/500) was added to each flask. Cells were incubated at 37° C, 5% CO2 for 4 days until extensive cytopathogenic effect was observed. Supernatant was harvested and clarified by centrifugation at 2000 rpm for 10 minutes in a benchtop centrifuge. Supernatant was aliquoted and frozen at -80°C.

### High-throughput live virus microneutralization assay

High-throughput live virus microneutralisation assays were performed as previously described ^14^. In brief, Vero E6 cells (Institut Pasteur) at 90-100% confluency were infected with given SARS-CoV-2 variants in 384-well format, in the presence of serial dilutions of patient serum samples. After infection, cells were fixed with 4% final Formaldehyde, permeabilized with 0.2% TritonX-100, 3% BSA in PBS (v/v), and stained for SARS-CoV-2 N protein using Biotin-labelled-CR3009 antibody produced in-house together with a Streptavidin-Alexa488 (S32354, Invitrogen) and cellular DNA using DAPI (10236276001, Merck). Whole-well imaging at 5x was carried out using an Opera Phenix (Perkin Elmer) and fluorescent areas and intensity calculated using the Phenix-associated software Harmony (Perkin Elmer). Inhibition was estimated from the measured area of infected cells/total area occupied by all cells and expressed as percentage of maximal (virus only wells). The inhibitory profile of each serum sample was estimated by fitting a 4-parameter dose response curve executed in SciPy. Neutralizing antibody titers are reported as the fold-dilution of serum required to inhibit 50% of viral replication (IC50), and are further annotated if they lie above the quantitative (complete inhibition) range, below the quantitative range but still within the qualitative range (i.e. partial inhibition is observed but a dose-response curve cannot be fit because it does not sufficiently span the IC50), or if they show no inhibition at all. Human coronavirus OC43 (HCoV-OC43) neutralization was performed as above, except Vero E6 cells were substituted for Mv1Lu cells.

### ELISA and other serological testing

Anti-S1 was performed as described previously ^32^. To minimize variation across ELISA plates, we re-scaled serum OD405 measurements by (i.) subtracting the *plate-wide* average negative control, (ii.) dividing by the *plate-wide* average positive control and then (iii.) multiplying by the *study-wide* median of the plate-averaged positive controls.

### Anti-nucleocapsid IgG detection

Anti-nucleocapsid IgG was measured using the Elecsys Anti-SARS-COV-2 assay (Roche; 09203095190) run on a Cobas e411 analyser (Roche) in accordance with the manufacturer’s instructions. Serum was used for this immunoassay and results reported as reactive (positive) or non-reactive (negative), with a semi-quantitative titer. To separate participants into anti-N positive and negative groups, we used their most recent anti-N result.

### Mass cytometry sample processing

Peripheral blood mononuclear cells were thawed in the presence of benzonase (Merck 70746-3, used at 1µl/ml), cells were counted and up to 3 x 10^6^ cells were processed for mass cytometry. Mass cytometry staining was performed using the MaxPar Direct Immune Profiling Assay (Fluidigm, now Standard Biotools), in line with the manufacturer’s instructions, with T cell expansion panel 3. Once stained and fixed, cells were stored at -80C and processed in batches. After thawing, cells were stained with Iridium as per the manufacturer’s instructions (Standard Biotools). Events were collected using a CyTOF XT (Standard Biotools), and were bead-normalized using the in-built algorithm.

| Target antigen | Clone | Isotope |
| --- | --- | --- |
| CD45 | HI30 | 89Y |
| CCR6 (CD196) | G034E3 | 141Pr |
| CD123 | 6H6 | 143Nd |
| CD19 | HIB19 | 144Nd |
| CD4 | RPA-T4 | 145Nd |
| CD8a | RPA-T8 | 146Nd |
| CD11c | Bu15 | 147Sm |
| CD16 | 3G8 | 148Nd |
| CD45RO | UCHL1 | 149Sm |
| CD45RA | HI100 | 150Nd |
| CD161 | HP-3G10 | 151Eu |
| CCR4 (CD194) | L291H4 | 152Sm |
| CD25 | BC96 | 153Eu |
| CD27 | O323 | 154Sm |
| CD57 | HNK-1 | 155Gd |
| CXCR3 (CD183) | G025H7 | 156Gd |
| CXCR5 (CD185) | J252D4 | 158Gd |
| CD28 | CD28.2 | 160Gd |
| CD38 | HB-7 | 161Dy |
| NCAM (CD56) | NCAM16.2 | 163Dy |
| TCRgd | B1 | 164Dy |
| CD294 | BM16 | 166Er |
| CCR7 (CD197) | G043H7 | 167Er |
| CD14 | 63D3 | 168Er |
| CD3 | UCHT1 | 170Er |
| CD20 | 2H7 | 171Yb |
| CD66b | G10F5 | 172Yb |
| HLA-DR | LN3 | 173Yb |
| IgD | IA6-2 | 174Yb |
| CD127 | A019D5 | 176Yb |
| Live/dead intercalator |  | 103Rh |
| DNA1 |  | 191Ir |
| DNA2 |  | 193Ir |
| CD134/OX40 | ACT35 | 142Nd |
| TIGIT | MBSA43 | 159Tb |
| CD69 | FN50 | 162Dy |
| CD279/PD-1 | EH12.2H7 | 165Ho |
| CD366/Tim-3 | F38-2E2 | 169Tm |
| CD278/ICOS | C398.4A | 175Lu |
| CD137/4-1BB | 4B4-1 | 209Bi |

### Mass cytometry gating strategy

Events from bead-normalized FCS files were gated as shown in Figs. S5 & S6. This was performed using R v4.0.0, the following packages: flowCore v2.2.0 ^33^, flowWorkspace v4.2.0, openCyto v2.2.0 ^34^ and ggcyto v1.18.0 ^35^.

Gated samples were re-saved as FCS files (this allowed the parallel processing of samples). Gated FCS files were analyzed using CATALYST ^36^. As a quality control step to further filter out dead cells, or debris, we performed clustering within CATALYST (which itself uses rounds of flowSOM ^37^ aggregated using ConsensusClusteringPlus ^38^), using the live/dead, DNA1 and DNA2 channels only (Fig. S7). This returned 3 meta-clusters, one of which was DNA1-and DNA2-negative, one was dead+ and the largest cluster contained DNA1+, DNA2+,live events. The DNA1+, DNA2+, live cluster was selected and used for all downstream analyses. PBMC were then re-clustered using CATALYST and the following markers (selected to define “types” of cells in PBMC): CD3, CD20, CD19, CD14, CD16, CD161, CD56, CD45RA, CD45RO, CD4, CD8a, CD11c, live/dead and DNA1 and DNA2. The PBMC dataset was sub-sampled to 500 cells/sample for visualization with UMAP-embedding and summary heatmap of marker expression. For differential abundance analyses, edgeR ^39^ was used via diffcyt ^40^. For differential state analyses, we used limma ^41^ via diffcyt. For each population of PBMCs, events were filtered and then re-clustered using CATALYST and differential expression analyses as before. The optimum number of meta-clusters was determined by visual inspection of UMAP projections and heatmaps for each cell type.

### Time to event analysis

Time to event analysis was performed in R using the survival package. Individuals were left censored on the day they received their third dose. Individuals were right censored on the date of an infection with a variant that was Omicron BA.2, at the last visit date (participants are asked if they have experienced COVID-19 symptoms in the interim, and an anti-N IgG level is tested), or at the date of dose 4. Omicron BA.2 infection was confirmed by viral sequencing, by S gene target presence, or if no nucleic acid testing was available, based upon calendar date. Infection >14d after dose 3 was considered the event of interest and included infections were the BA.1/BA.2 BA.2-BA.4/5 assignment was date-based. Days of entry, exit and event were calculated with respect to the earliest date for dose 3 in the study. Data are presented as cumulative incidence plots, with at numbers at risk shown. Cox proportional hazard models were used as described in the text.

### Data analysis

Study data were collected and managed using REDCap electronic data capture tools hosted at University College London ^42, 43^. Data were exported from REDCap into R for visualization and analysis, similar to previously ^16^. Neutralizing antibody titers are reported as IC_50_ values. As described above, for each serum sample, four dilutions (1:40, 1:160, 1:640, 1:2560) are assayed in duplicate. All 8 points are used to fit a 4 parameter curve, and the IC_50_ (the fold-dilution corresponding to 50% viral inhibition), is reported. IC_50_ values below 40 and above 2560 are reported as ‘weak’ or ‘complete’ inhibition. For plotting and analysis, winsorizing was used: IC_50_ values above the quantitative limit of detection of the assay (>2560) were recoded as 5120; IC50 values below the quantitative limit of the assay (<40) but within the qualitative range were recoded as 10; data below the qualitative range (i.e. no response observed) were recoded as 5.

All data analysis was performed in R. The statistical tests used are described in the relevant section of the methods, figure legends or text.

### Online supplemental material

Fig S1 shows receiver operating characteristics for anti-S1 and anti-N IgG for the prediction of serological profile. Fig S2 displays longitudinal PCRs, symptom diaries and anti-N IgG to confirm seronaive individuals. Fig S3 shows the demographics, vaccine usage and dosing intervals for broad anti-N IgG seropositive or seronegative individuals. Fig S4 shows the neutralization of HCoV-OC43 is not augmented by SARS-CoV-2 Spike exposure (infections or vaccinations). Figure S5-S7 show the mass cytometry gating strategy, illustrative gating and quality control. Figure S8 shows PBMC-level analysis of mass cytometry.

### Data availability

Requests for de-anonymized data will be considered by the Legacy Governance Board, via, to ensure the request is from a genuine researcher and that legal and ethical obligations are maintained.

## Acknowledgements

This work was supported by the National Institute for Health Research University College London Hospitals Department of Health’s NIHR Biomedical Research Centre (BRC), as well as by the UK Research and Innovation and the UK Medical Research Council (MR/W005611/1 [Genotype to Phenotype consortium], Rosetrees [M926]), and by the Francis Crick Institute which receives its core funding from Cancer Research UK (CC1283, CC2230, CC2166, CC2087, CC2060, CC2041, CC2112), the UK Medical Research Council (CC1283, CC2230, CC2166, CC2087, CC2060, CC2041, CC2112), and the Wellcome Trust (CC1283, CC2230, CC2166, CC2087, CC2060, CC2041, CC2112).

The authors would like to thank all the study participants, the staff of the NIHR Clinical Research Facility at UCLH including Kirsty Adams and Marivic Ricamara. We would like to thank Jules Marczak, Gita Mistry, Simon Caiden, Matala Dyke, and the staff of the Scientific Technology Platforms (STPs) and COVID-19 testing pipeline at the Francis Crick Institute. We thank Prof. Wendy Barclay of Imperial College and the wider Genotype to Phenotype consortium for the Alpha and Delta strains used in this study, and Max Whiteley and Thushan I de Silva at The University of Sheffield and Sheffield Teaching Hospitals NHS Foundation Trust for providing source material. We thank Prof. Gavin Screaton of the University of Oxford for the Omicron BA.1 strain used in this study. We thank Khadija Kahn and Alex Sigal of the Africa Health Research Institute for the Beta and Omicron BA.5 strain used in the study, and Ann-Katrin Reuschl and Clare Jolly of University College London for propagation. We thank Dr Laura McCoy of UCL for her original synthesis of the CR3009 protein used in development of the HTS assay. We also thank Marg Crawford, Robert Goldstone, and Harshil Patel for generation and processing of sequencing data. We are grateful to the Crick’s Legacy Immunology Advisory Group (chaired by Prof. Carola Vinuesa FRS) for their ongoing support, and to Dr Laurie Tomlinson at London School of Hygiene and Tropical Medicine for helpful discussions. This research was funded in whole, or in part, by the Wellcome Trust (CC1283, CC2230, CC2166, CC2087, CC2060, CC2041, CC2112). For the purpose of Open Access, the author has applied a CC-BY public copyright licence to any Author Accepted Manuscript version arising from this submission.

## Author contributions

EJC conceptualized the project, performed the analysis, co-wrote the draft and edited the final manuscript. HT performed clinical metadata curation and laboratory data generation. MYW, KAW, PSH, DL, SN, SH, HVM, AH, M., LSH, ASH, BC and MM generated laboratory data and processed blood samples. MYW and RH provided high-throughput live-virus microneutralisation expertise. EM and YN obtained ethical approval for the Legacy study, with supervision from CSw and SG. CS and GK performed data curation. VM, BZ and SJWE provided expert input for the time-to-event analyses. VL, AR, JN, NO’R, BW and MH provided supervision to teams generating and curating data. RCLB, DLVB and ECW co-wrote the manuscript. Funding was secured by EJC, CSw, SG, RCLB, DLVB, ECW led the clinical and laboratory teams to generate the data presented here. EJC, RCLB, DLVB and ECW are senior authors.

---

**Table S1.** Demographics and vaccine characteristics of the mass cytometry cohort. The primary doses, third doses, age, sex and interval between dose 2 and 3, and cumulative anti-N IgG status are summarized for individuals in the mass cytometry dataset

| Characteristic | **broad, N = 11<sup>1</sup>** | **narrow, N = 6<sup>1</sup>** |
| --- | --- | --- |
| **doses 1 and 2** |  |  |
| AZD1222 | 0 (0%) | 0 (0%) |
| BNT162b2 | 11 (100%) | 6 (100%) |
| mRNA1272 | 0 (0%) | 0 (0%) |
| others | 0 (0%) | 0 (0%) |
| **dose 3** |  |  |
| AZD1222 | 0 (0%) | 0 (0%) |
| BNT162b2 | 11 (100%) | 6 (100%) |
| mRNA1272 | 0 (0%) | 0 (0%) |
| others | 0 (0%) | 0 (0%) |
| **Sex** |  |  |
| Female | 9 (82%) | 3 (50%) |
| Male | 2 (18%) | 3 (50%) |
| $\chi^2$ test $P=0.4$ | | |
| **Age** | 56 [54-61] | 62 [48-64] |
| 2 tailed Mann Whitney $P=0.6$ | | |
| **Interval between dose 3 and 2 [days]** | 193.0 [188.5-195.0] | 192.5 [189.0-198.2] |
| 2 tailed Mann Whitney $P=0.6$ | | |
| **anti-N IgG results up-to 28d after dose 3** |  |  |
| all negative | 11 (100%) | 6 (100%) |

<sup>1</sup>n (%); Median [25%-75%]

**Figure S1.**
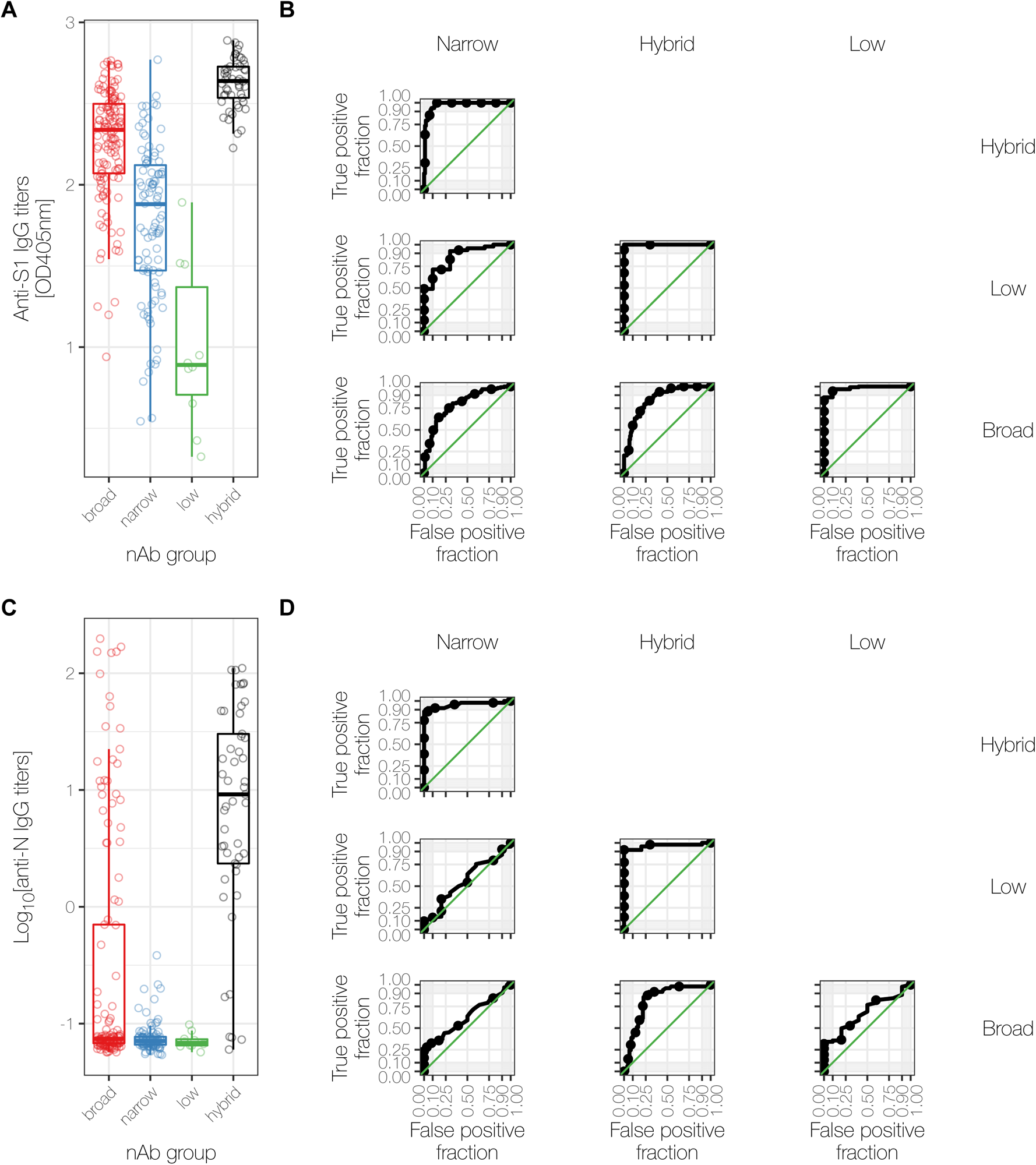
Receiver operating characteristic for anti-S1 and anti-N IgG predicting serological profile. **(A)** Titres of anti-S1 IgG, reported as scaled absorbance at OD405nm, for all 4 serological profile groups. **(B)** Receiver operating characteristic (ROC) curves for anti-S1 IgG titres in (A), between the indicated serological profile groups. **(C)** Decimal logarithm of anti-N IgG titres for all 4 serological groups. **(D)** As in (B), using anti-N IgG titres from (C).

**Figure S2.**
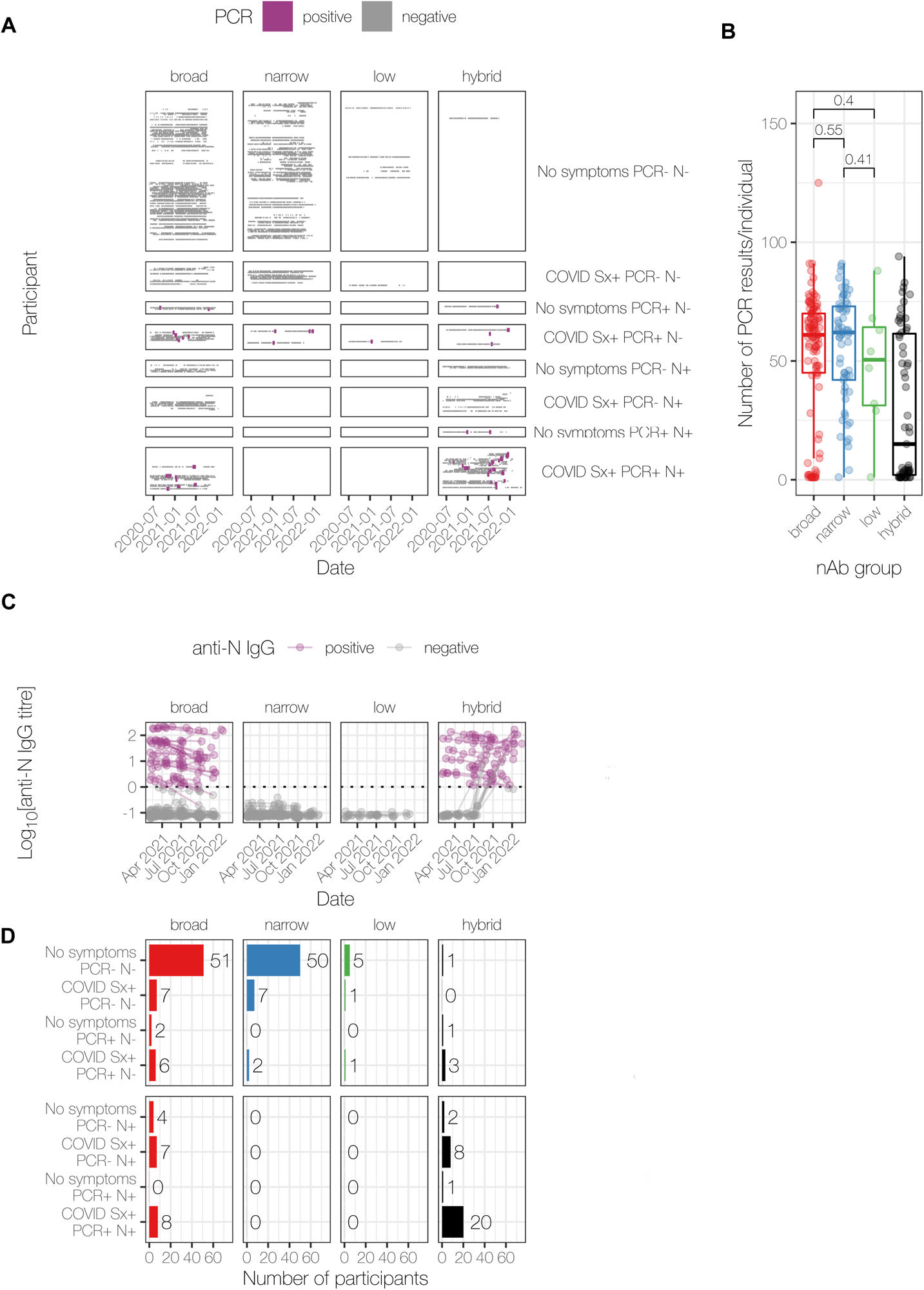
Longitudinal PCR screening, symptom diaries and anti-N IgG titres confirm seronaïve individuals. **(A)** Longitudinal PCR screening results are shown for each individual (as a row) over time (date), separated by the 4 serological profiles, and by the reporting of prior symptoms, or by the presence of anti-N IgG (at any time up to dose 3). **(B)** The number of PCR tests per individual is shown for each serological profile. **(C)** Longitudinal anti-N IgG titres are shown for each of the 4 serological groups, up to dose 3. No broad responders gained anti-N IgG during the course shown **(D)** The number of participants within the following groups: Symp toms (No symptoms or COVID Sx+), PCR (PCR- or PCR+), anti-N IgG (N+ or N-), stratified by serological groups, using syptom diaries, PCR testing and anti-N titers up to dose 3. In (B), P values from unpaired two tailed Mann-Whitney tests are shown. The hybrid group has lower testing since individuals were granted a grace period before re-commencing screening after a positive test.

**Figure S3.**
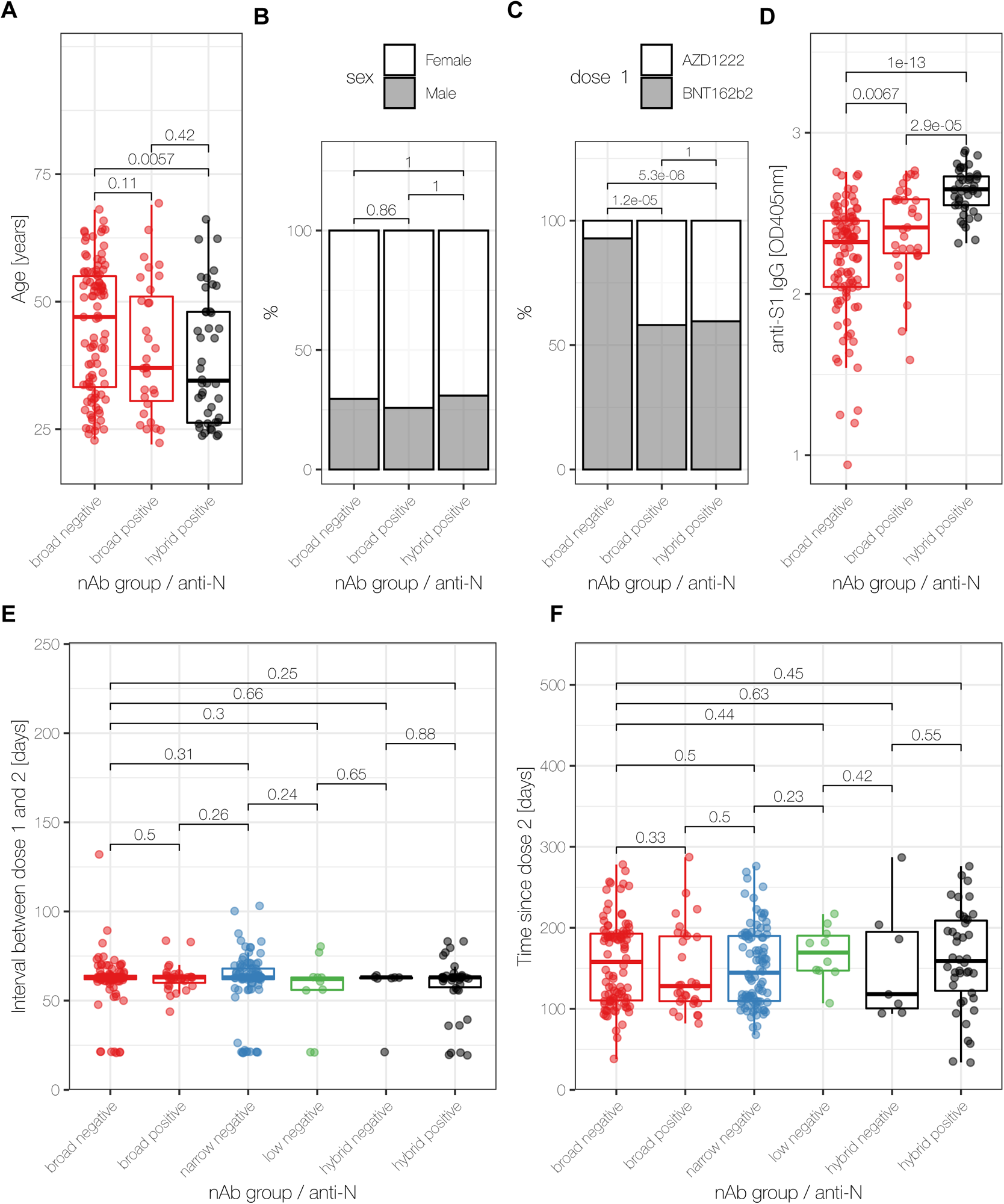
Demographics, vaccine usage and dose intervals for broad anti-N IgG seropositive or seronegative individuals. **(A)-(D)** Age in years (A) or sex (B) or vaccine used for doses 1 and 2 (C) or anti-S1 binding titres (D) for broad anti-N seropositive and seronegative individuals, compared to hybrid responders. **(E)-(F)** Time interval in days between doses 1 and 2 (E) or serum sampling time from dose 2 (F) for all serological groups, stratified by anti-N result. In (A, D-F), P values shown are from two tailed unpaired Mann-Whitney tests, without multiple correction testing. In (B-C) P values are from χ^2^ tests.

**Figure S4.**
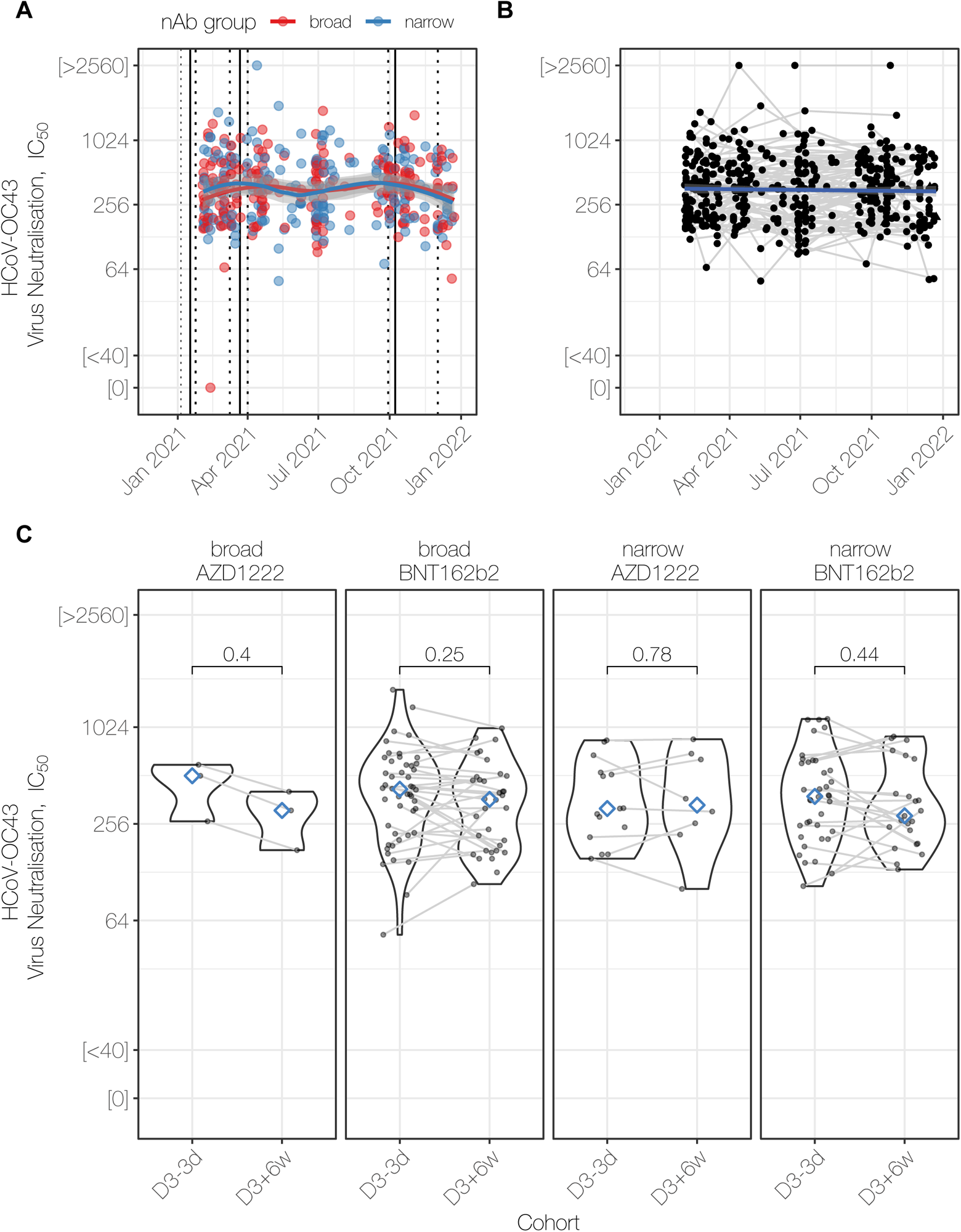
Neutralization of the seasonal human coronavirus HCoV-OC43 is not augmented by SARS-CoV-2 vaccination or infection. **(A)** HCoV-OC43 live-virus microneutralization titer trajectories for broad and narrow responders. The median dates of vaccine doses and their interquartile ranges are shown by the vertical solid and dashed lines respectively. Neutralization titers are expressed as reciprocal of dilution with 50% inhibition of viral infection (IC_50_). **(B)** As in (A), with a linear regression fit to demonstrate rate of waning **(C)** HCoV-OC43 neutralization before (median -91d; IQR 9-77d) and up to 6 weeks after (median 23d, IQR 18-31d) dose 3 in broad and narrow responders, stratified by their primary vaccination course.

**Figure S5.**
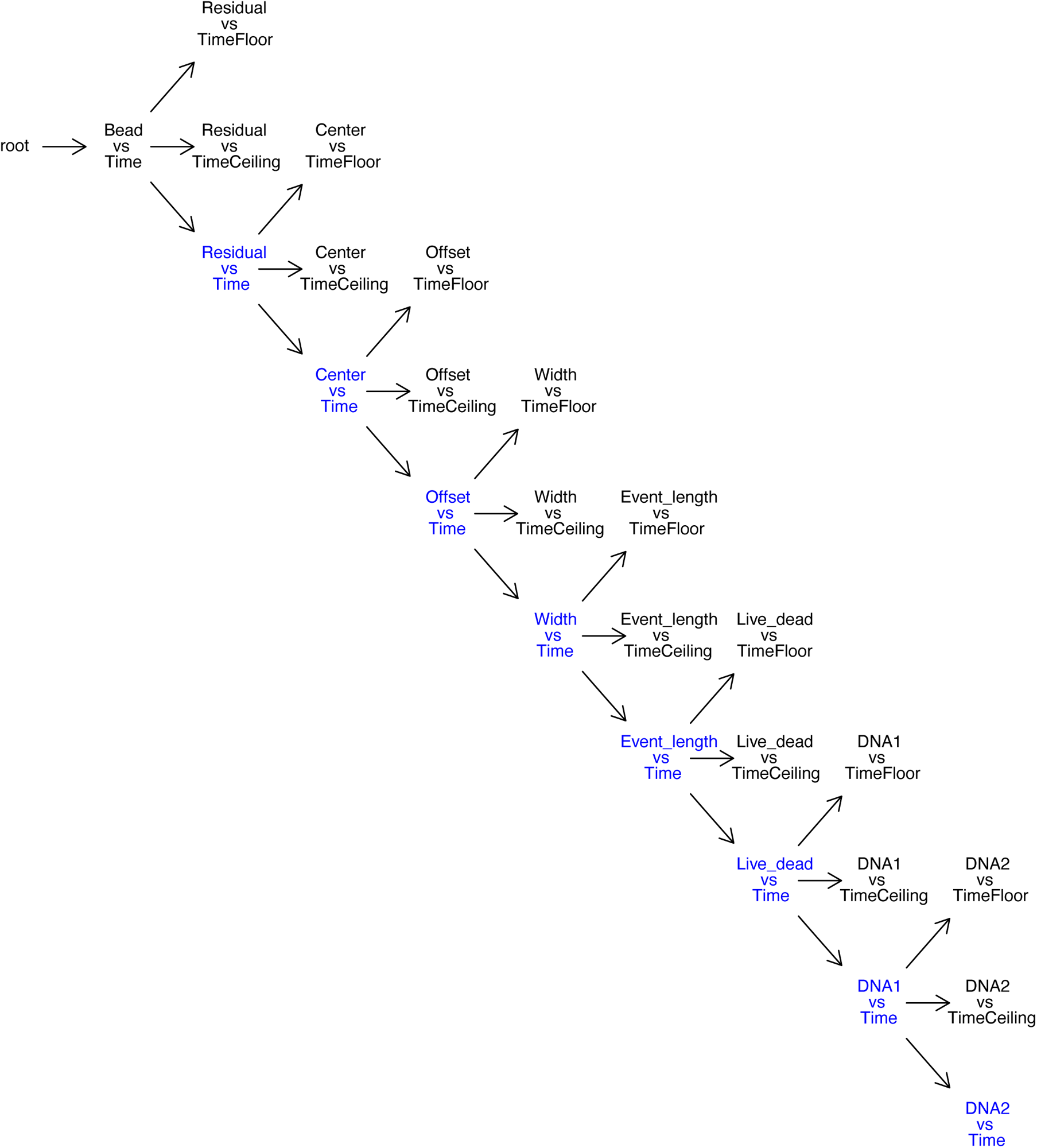
Mass Cytometry gating tree. Gating hierarchy progressing from raw events to processed single cells for downstream analysis. See Figure S6 for illustrative gates, and Figure S7 for flowSOM based quality control before biological clustering.

**Figure S6.**
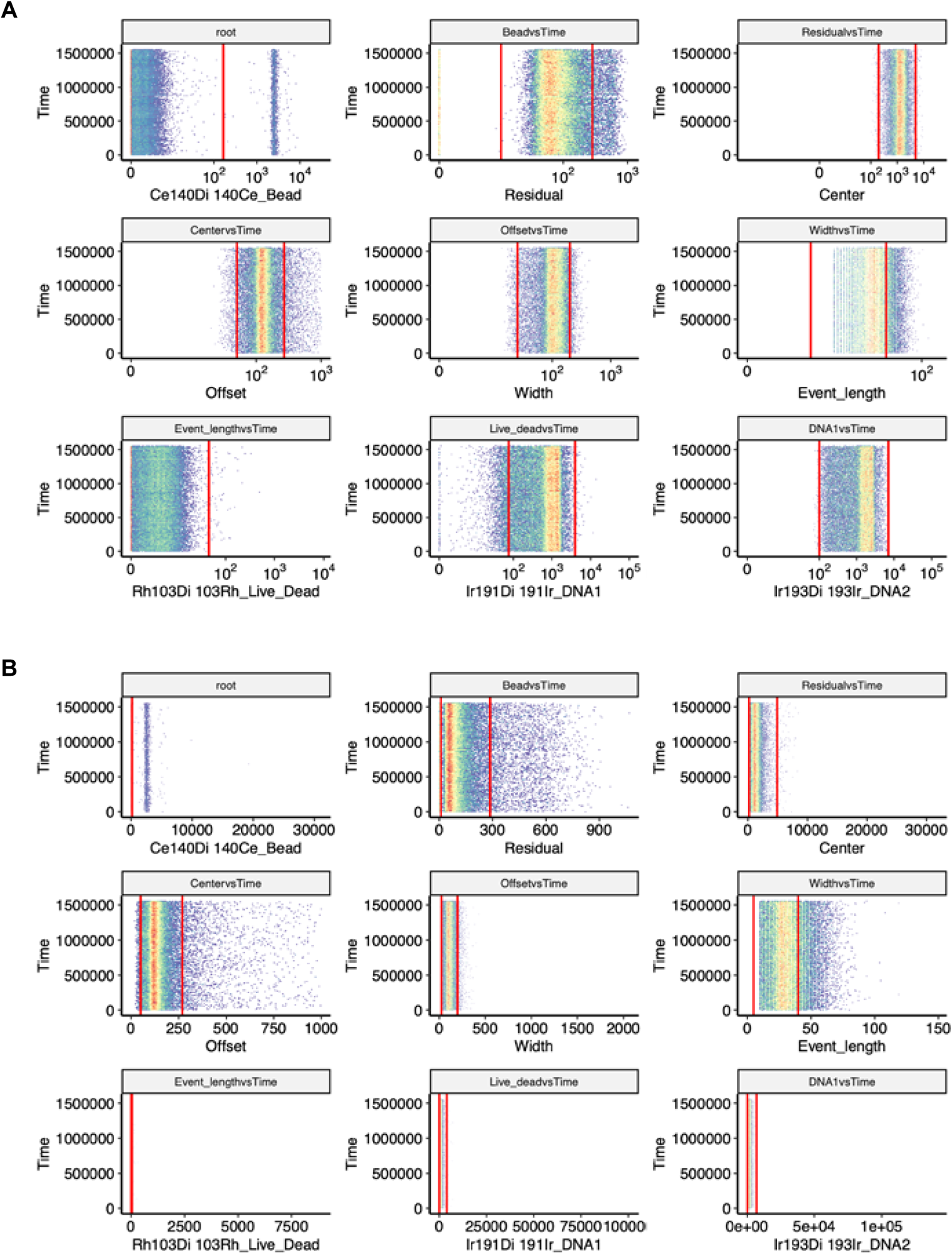
Mass Cytometry gating strategy. **(A)** and **(B)** Sequential gating of events. First QC-beads are gated out (top left) and samples progress through the gating hierarchy (Figure S5), by row left-right to processed single cells. In (A) parameters are plotted after inverse hyperbolic sine transformation (fasinh) and in (B) the same parameters are plotted as a linear transform. Details of the gating algorithm are described in the Methods.

**Figure S7.**
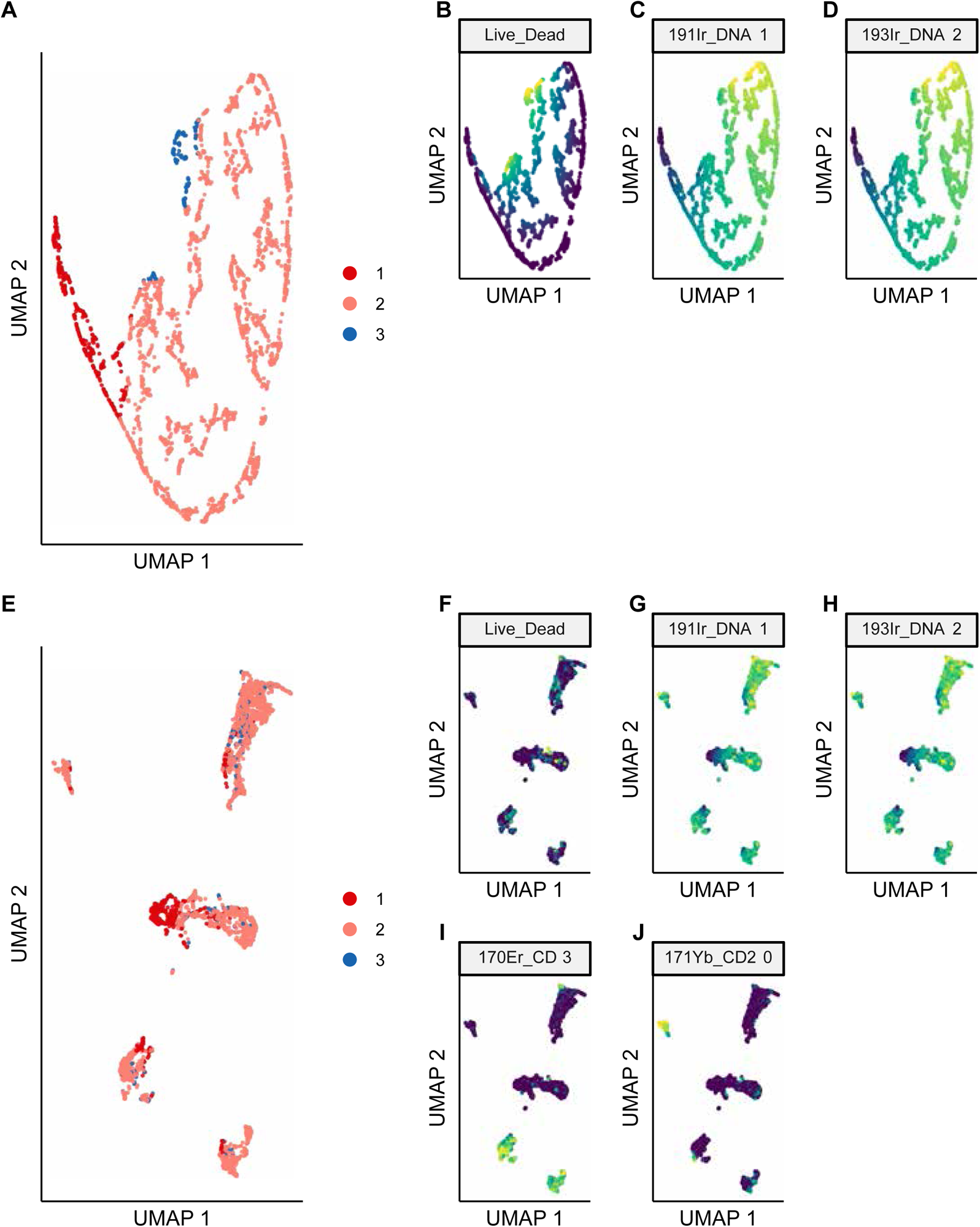
Mass cytometry quality control using live/dead and DNA1 + DNA2. **(A)** UMAP embedding of gated events, after flowSOM and ConsensusClusterPlus performed with the following channels: live/dead exclusion, DNA1 and DNA2. **(B)-(D)** UMAP embedding from (A) shaded by the intensity of signal from each channel: live/dead exclusion (B), DNA1 (C) and DNA2 (D). **(E)** UMAP embedding of gated events, after flowSOM and ConsensusClusterPlus performed with the following channels: CD3, CD20, CD19, CD14, CD16, CD161, CD56, CD45RA, CD45RO, CD4, CD8a, CD11c, live/dead and DNA1 and DNA2. The shading reflects the clustering in (A). **(F)-(J)** UMAP embedding from (E) shaded by the indicated marker.

**Figure S8.**
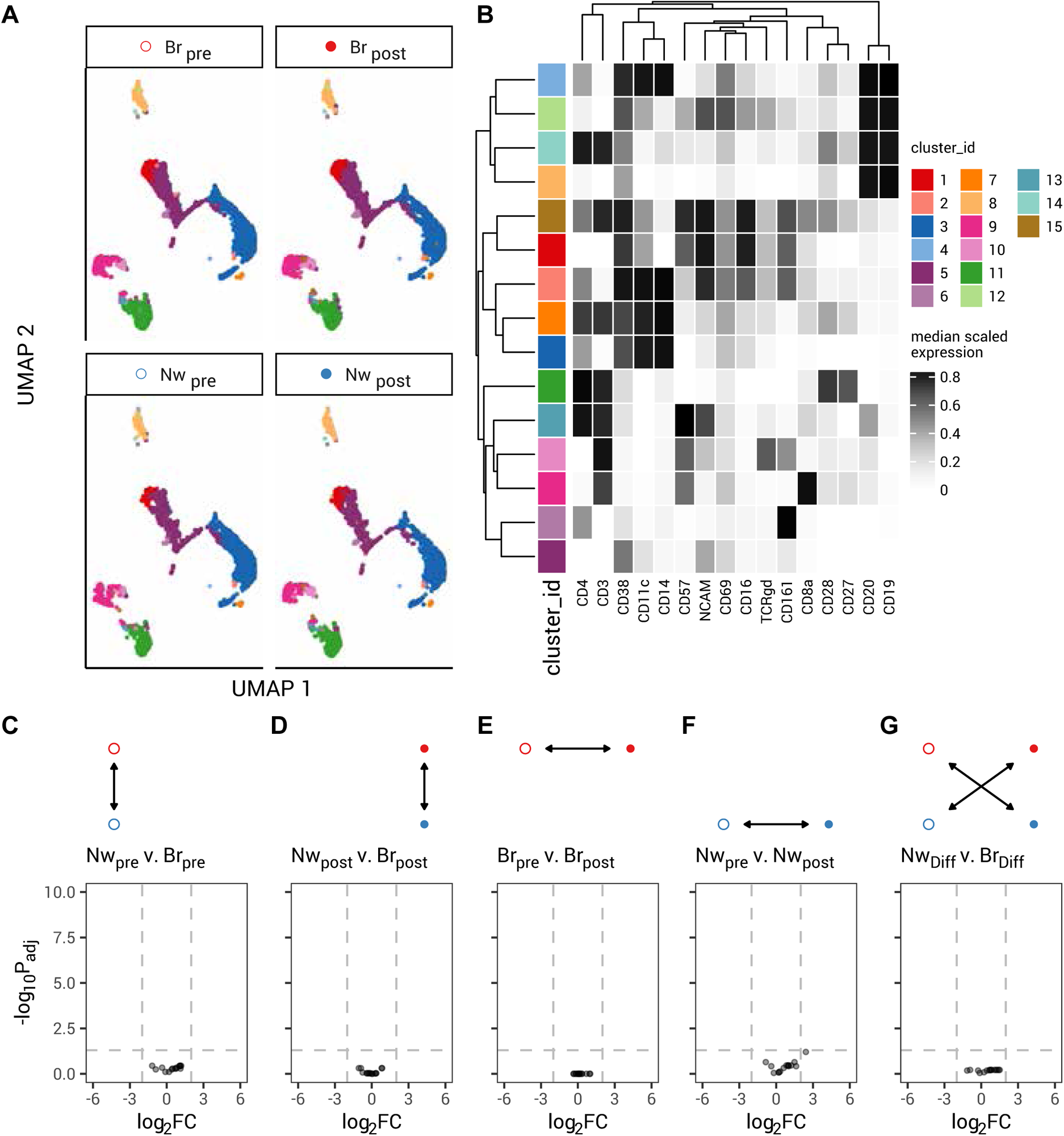
PBMC-level mass cytometry analysis. **(A)** UMAP embedding of peripheral blood mononuclear cells (PBMC) separated by breadth (Nw narrow, n=6; Br broad, n=11) and before (pre) and after (post) vaccination. 15 clusters identified by FlowSOM and ConsensusClusteringPlus are shaded. **(B)** Heatmap of surface expression of selected markers for the clusters shown in (A). Rows represent the clusters shown in (A), and their color key is shared. Columns reflect the labelled cell surface marker. Scaled expression is shown from blue (low/no expression) to yellow (high expression). Cluster 9 are CD8^+^ T cells, with the adjacent cluster 10 being TCRγδ^+^ T cells. **(C)-(G)** Differential abundance analysis for the 15 PBMC clusters shown in (A) and (B), for the comparisons indicated: narrow pre vs broad pre in (C); narrow post vs broad post in (D); broad pre vs.broad post in (E); narrow pre vs.narrow post in (F); and the difference between (narrow pre vs. narrow post) and (broad pre vs. broad post) in (G). log_2_ fold change ±1 and *P*_adj_=0.05 are shown by dashed lines.

